# Preneoplastic cells switch to Warburg metabolism from their inception exposing multiple vulnerabilities for targeted elimination

**DOI:** 10.1101/2023.01.09.523333

**Authors:** Henna Myllymäki, Lisa Kelly, Abigail M. Elliot, Roderick N. Carter, Nicholas M. Morton, Yi Feng

## Abstract

Otto Warburg first described tumour cells as displaying enhanced aerobic glycolysis whilst maintaining defective oxidative phosphorylation (OXPHOS) for energy production almost 100 years ago ^1,2^. Since then, the ‘Warburg effect’ has been widely accepted as a key feature of rapidly proliferating cancer cells^3,4^. Targeting cancer metabolism is now being considered as a promising precision oncology therapeutic approach^5^. What is not clear is how early “Warburg metabolism” initiates during cancer progression and whether changes in energy metabolism might influence tumour progression *ab initio*. We set out to investigate energy metabolism in HRAS^G12V^ driven preneoplastic cell (PNC) at inception, in a zebrafish skin PNC model; and to test how the impact of manipulating energy metabolism in the whole animal may impact PNC initiation. We find that, within 24 hours of HRAS^G12V^ induction, PNCs upregulate “Warburg metabolism”, and that this is required for their expansion. We show that blocking glycolysis reduces PNC proliferation, whilst increasing available glucose both enhances PNC proliferation and also reduces apoptosis. Impaired OXPHOS of PNCs might be exploited therapeutically since a mild complex I inhibitor, metformin, selectively induces apoptosis and suppresses proliferation of PNCs. In addition, we find mitochondrial fragmentation in PNCs occur prior to metabolic alteration and this is important for their survival since exposure to Drp1/Dnml1 inhibitor, mdivi, which blocks mitochondrial fragmentation leads to enhanced PNC apoptosis. Our data indicate that altered energy metabolism is one of the earliest events upon oncogene activation in somatic cells, which provide a targeted and effective tumour prevention therapy.

**Key findings:**

- Glycolysis is upregulated in HRAS^G12V^ expressing preneoplastic cells (PNCs) in zebrafish skin, and is required for PNC proliferation
- OXPHOS respiration is impaired in PNCs and metformin complex I inhibition specifically eliminates PNCs
- PNC undergo mitochondrial fragmentation and exhibit reduced membrane potential
- Mdivi reverses mitochondrial fragmentation in PNCs and triggers apoptosis

## Introduction

Somatic cells that acquire an oncogenic mutation have an ability to undergo expansion and form pre-neoplastic lesions in vivo. However, little is known about the mechanisms involved in driving their initial clonal expansion^6^. Altered metabolism is one of the hallmarks of cancer, enabling cancer cells to sustain uncontrolled growth even under restricted nutrient conditions^7^. Amongst metabolic changes in cancer cells, the “Warburg Effect” is the most well-known and characterized by increased uptake of glucose and its conversion to lactate by glycolysis under normoxia conditions^4^. Although inefficient for ATP generation, Warburg metabolism provides advantages to fast proliferating cells, to meet their bio-synthetic demand ^4^. Warburg metabolism has been exploited to develop targeted cancer therapeutics and diagnostic tools, however, its role in the earliest stage of tumourigenesis is less clear. Altered mitochondrial morphology and function are crucial for cancer cell survival and tumourigenesis in several models^8^. Dysregulation of proteins that modulate mitochondrial dynamics is a targetable feature in human tumours and there is increasing interest in the development of novel cancer therapies by targeting the mitochondrion^9^. However, most studies on the susceptibility to metabolic targeting have employed established tumours, and little is known of the initial stages of tumour development and what are the metabolic features of the pre-neoplastic cell. Could early metabolic changes at the pre-neoplastic stage offer a novel Achilles heel for therapeutic chemoprevention? The lack of appropriate model systems has hindered our understanding of the initial events in vivo during pre-neoplastic development. Here we use a zebrafish skin pre-neoplastic cell (PNC) model, which focuses on a common oncogenic mutation, the human hRAS^G12V^, to investigate alterations in cellular energy metabolism at the initial stage of tumourigenesis and to ask whether this can be targeted for PNC elimination.

## Results and Discussion

### 1. Glycolysis is enhanced in PNCs to boost their proliferation and supplemental glucose promotes PNC expansion

In our model of tumour initiation, human HRAS^G12V^ (thereafter HRAS) was induced in zebrafish larval keratinocytes to mimic the initial mutational event leading to PNC development (Figure 1 A Schematic). EGFP tagged PNCs can be identified from 8 hours post induction (hpi) and analysing cell proliferation suggests that enhanced PNC growth is overt from 24hpi and the proliferation rate in PNCs remains high at later times^10,11^. We therefore focused our analysis within the first 24 hours of PNC emergence to establish the earliest metabolic changes following PNC induction.

**Figure 1.**
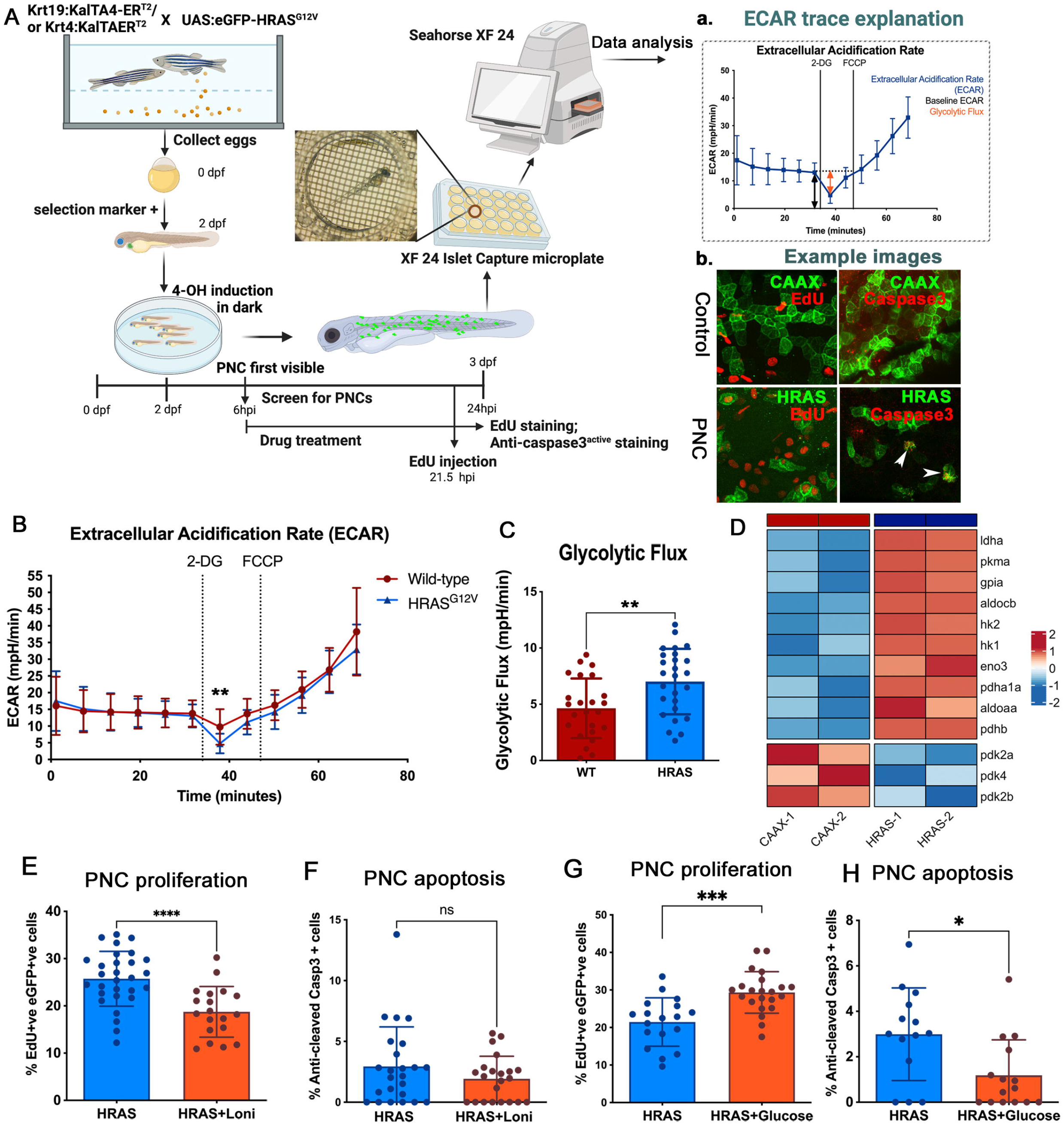
Glycolysis is important in boosting PNC proliferation and excess glucose promotes PNC survival and expansion. A. Schematic showing the inducible human HRAS^G12V^ mediated preneoplastic cell in zebrafish larval skin tissue. Seahorse^®^ Metabolic Flux analyses were carried out at 24hours post induction (hpi). A a, explanation of ECAR trace b, representative confocal images of EdU staining (proliferation) and anti-cleaved-Caspase3 staining (apoptosis) that were carried out in this study. B. Seahorse Analyser® ECAR readout over time, showing no difference in baseline ECAR (before cycle 6), glycolytic flux (after adding 2-DG) readout at cycle 7 showing enhanced glycolytic flux in HRAS PNCs. Note: 2-DG leads to a transient ECAR change which recovers from cycle 8, this is thought to be due to the whole organism response to 2-DG. FCCP was added (black line) to assess respiratory function using the complimentary OCR readout (data not shown, as similar OCR data were presented in figure 2). C. Quantification (cycle 7 ECAR) showing PNC have enhanced glycolytic flux (mean≥ SD, n≥ 24, p=0.0042). Hexokinase inhibitor lonidamine (2nM) treatment leads to reduced EdU positive cells in PNCs (unpaired t test, mean +/− SD, n≥19, p=0.0001). D. Pseudobulk differential expression analysis of single-cell RNA sequencing data, showing significantly up- and down-regulated genes related to glycolysis in HRAS PNCs vs. Control CAAX keratinocytes at 24 hpi. Heatmap depicts log fold-change (EdgeR, n=2, FDR < 0.05). E. Hexokinase inhibitor lonidamine (2nM) treatment did not change PNC apoptosis (unpaired t test, mean +/− SD, n≥ 22, p= 0.2042). F. Glucose injected larvae show increased PNC proliferation (unpaired t test, mean +/− SD, n≥ 18, p=0.0002). G Glucose injected larvae show decreased PNC apoptosis (unpaired t test, mean +/− SD, n≥ 14, p=0.0119).

In order to assess glycolytic activity of PNCs in skin tissue, we established a protocol using intact zebrafish larvae and the Seahorse analyzer^®^ to measure extracellular acidification rate (ECAR) (Figure 1 A schematic), which has generally been used as a proxy to evaluate aerobic glycolysis of cells and tissues (figure A a)^12^. In this assay, 2-Deoxy-d-glucose (2-DG) was used to block glycolysis generated ECAR and here we saw greater reduction of ECAR in PNC-bearing larvae compared with controls, indicating a higher rate of glycolysis in PNCs (figure 1 A a, B, C). Furthermore, we analysed gene expression of enzymes involved in the glycolysis pathway using existing single cell RNA sequencing data of 24hpi HRAS expressing PNCs (Elliot et al in preparation). We saw up-regulation of many key glycolytic enzymes including hexokinase 1 and 2 (*hk1, hk2*) and lactate dehydrogenase a (*ldha*) (figure 1D), thus further supporting enhanced glycolysis in PNCs.

To test whether enhanced glucose usage is required for PNC progression, we treated larvae with low doses of glycolysis inhibitor, Lonidamine (2nM)^13^, and this led to a significant reduction in PNC proliferation (figure 1 E), while the same dose had no impact on control EGFP-CAAX (CAAX thereafter) expressing keratinocyte proliferation (Supplement figure 1). We saw no significant increase in apoptotic cell death (figure 1 F), suggesting that glycolysis in PNCs was important for their proliferation but not their survival.

To further test the impact of altered energy availability, we mimicked the condition of surplus sugar availability by injecting glucose into the yolk (the energy storage tissue for fish larvae). This led to a marked enhancement of PNC proliferation (figure 1 G). Interestingly, apoptosis was also reduced in PNCs from larvae receiving a glucose supplement (figure 1 H). These data established a promotional role for glycolysis in PNC growth and provided direct evidence that increased energy supply in the form of glucose promotes cells with oncogenic potential to survive and expand in vivo.

### 2. OXPHOS Respiration is also altered in PNCs and targeting complex I with metformin can specifically eliminate PNCs in vivo

It is believed that enhanced glycolysis in cancer cells promotes the generation of building blocks required for the increased demands of cell mass growth, while in parallel OXPHOS is still required or may even be up-regulated to provide sufficient energy supply^14,15,16,17^. To assess mitochondrial respiratory activity of PNCs, we developed a protocol similar to the ECAR measurement, using the Seahorse Analyzer^®^ to measure Oxygen Consumption Rate (OCR) from intact larvae (Supplement figure 2). Mitochondrial un-coupler FCCP (Carbonyl cyanide-p-trifluoromethoxyphenylhydrazone) was used to allow us to measure mitochondrial maximum respiration rate (figure 2A). These data showed that whilst baseline respiratory capacity was not altered in larvae with PNCs, their maximum respiratory capacity was significantly decreased compared to CAAX controls (figure 2 A, B, C), indicating impaired mitochondrial function. Previous studies have suggested that cells with HRAS mutations in vitro develop complex I deficiency through rapid accumulation of mutations in complex I components, leading to reduced mitochondrial respiratory function^18^. Whilst we were unable to perform analyses to determine whether there were mutations or not in Complex I components of PNCs in vivo, we did observe a reduced capacity for respiratory function. Additionally, gene expression analysis revealed that OXPHOS genes were enriched in PNCs compared with CAAX controls (figure 2 C). We speculate that this could be a compensatory up-regulation of mitochondrial gene expression due to damaged mitochondria, which again might explain impaired mitochondrial function. We reasoned that cells with complex I deficiency would be more vulnerable to complex I perturbation, and therefore targeting complex I by biguanides could specifically eliminate PNCs versus healthy epithelia^19,20^. Biguanides are widely used as anti-hyperglycaemic agents that target complex I^21^ and have previously been used to target cancer cells carrying complex I deficiency^22^. Metformin is one of the most frequently used biguanides for diabetes mellitus and prediabetes treatment^21^. Interestingly, previous reports have suggested that metformin leads to reduced risk of Oesophageal Squamous Cell Carcinoma^23^ and metformin was shown to reduce tumourigenesis through inhibiting complex I in xenograft models^24^. Due to established correlation that metformin medication reduced cancer in diabetic patients and positive outcome from pre-clinical studies of anti-cancer function of biguanides, there is an interest in repurposing metformin for cancer therapy with multiple clinical trials ongoing ^25,26^. We treated PNC – bearing larvae with metformin for 4 hours and this leads to increased activated-caspase-3 staining in PNCs in the superficial skin cell layer (figure 1 D). A longer treatment of 8 hours also led to a significant increase of activated caspase-3 signal in basal PNCs (figure 2 E). Furthermore, we detected reduced EdU incorporation in PNCs compared with untreated controls (figure 4 B) indicating that OXPHOS is important for PNC proliferation and that targeting complex I with metformin can induce PNC apoptosis. Importantly, metformin treatment had no effect on healthy skin cells (Supplement figure 2) and there were also no visible adverse effects on larval survival. Therefore, we have established that metformin, through targeting complex I, can eliminate HRAS mediated PNCs in vivo at inception and provide further evidence for potent cancer preventive efficacy of metformin.

**Figure 2.**
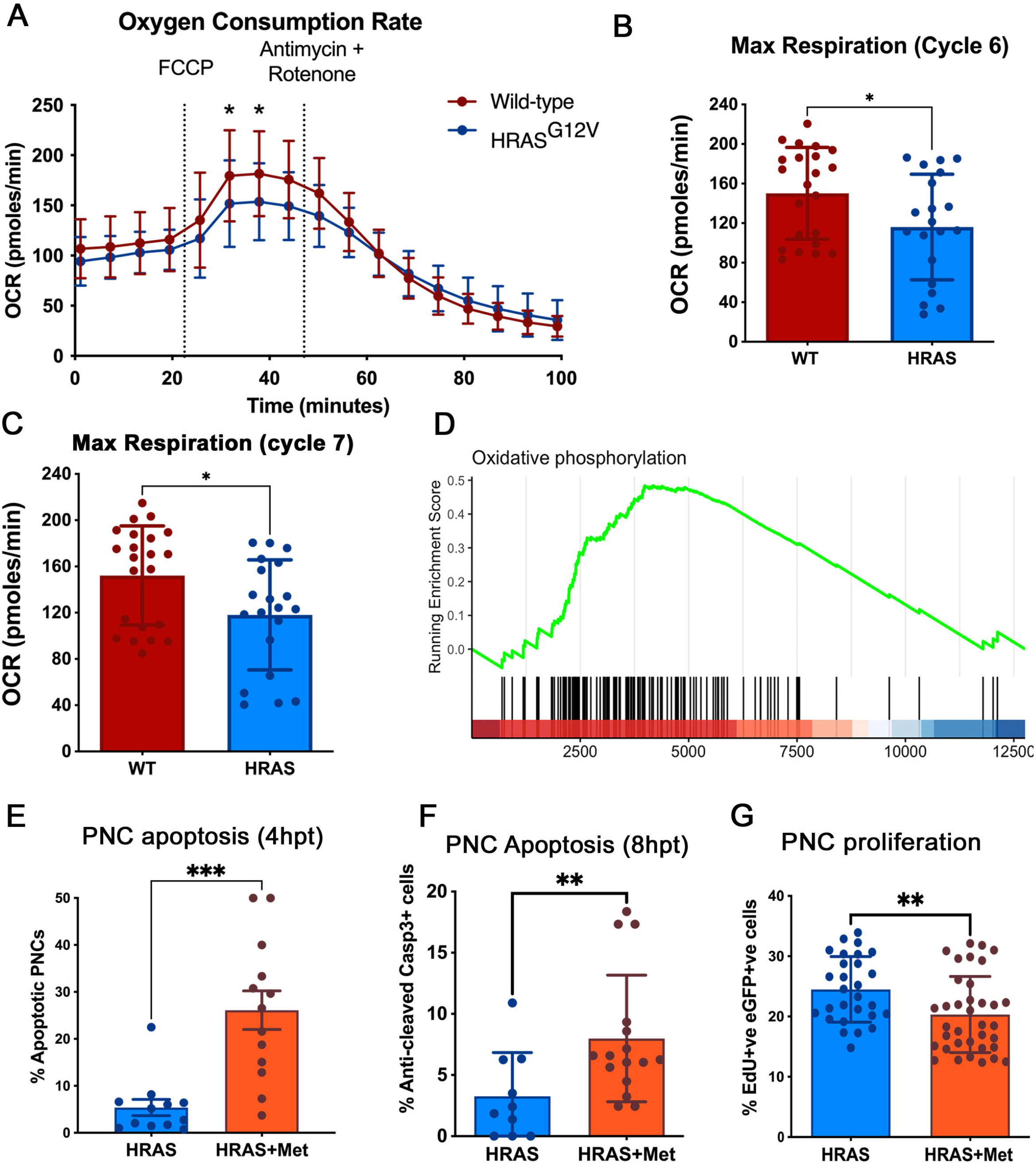
OXPHOS is impaired in PNCs and complex I inhibitor metformin suppresses PNC proliferation and induces PNC apoptosis in vivo. A. Seahorse Analyser^®^ Oxygen Consumption Rate (OCR) measurement comparing control larvae with PNC bearing larvae, graph showing OCR readout over time. Cycle 4 showing similar baseline respiration. Uncoupler FCCP was added after cycle 4 (black line) which assesses the reserve OCR. Cycle 7 showing significantly reduced maximum OCR in PNC bearing larvae, indicating reduced reserved respiration capacity. After cycle 8, treatment with the complex III inhibitor antimycin and the complex I inhibitor rotenone (black line) allowed the non-respiratory contribution to OCR to be determined, and there was no difference detected. (p< Mean +/− SD, n ≥ 20 embryos, Two-way ANOVA followed by Sidak’s multiple comparisons test.) B. Quantification showing maximum respiratory capacity is reduced in HRAS expressing PNCs (cycle 6 OCR, unpaired t test, mean+/−SD, n≥20, p=0.0326). C. Quantification showing maximum respiratory capacity is reduced in HRAS expressing PNCs (cycle 7 OCR, mean +/− SD unpaired t test, n≥20, * p= 0.0193) D. Gene-set enrichment analysis of single-cell RNA sequencing data shows that oxidative phosphorylation is enriched in HRAS expressing PNCs vs. CAAX expressing control keratinocytes (NES= 1.4401, p= 0.0409, FDR= 0.1308). E. Quantification showing metformin treatment induces superficial skin PNC apoptosis within 4hpt (Mean+/− SEM, Mann-Whitney test, n≥12, p=0.0001). F. Quantification showing metformin treatment induces basal skin PNC apoptosis (Mean+/− SD, unpaired t test, n≥10, p=0.0185). G. quantification showing reduced proliferation of basal PNCs upon metformin treatment (Mean+/− SD, unpaired t test, n≥ 28, p=0.0072).

**Figure 3.**
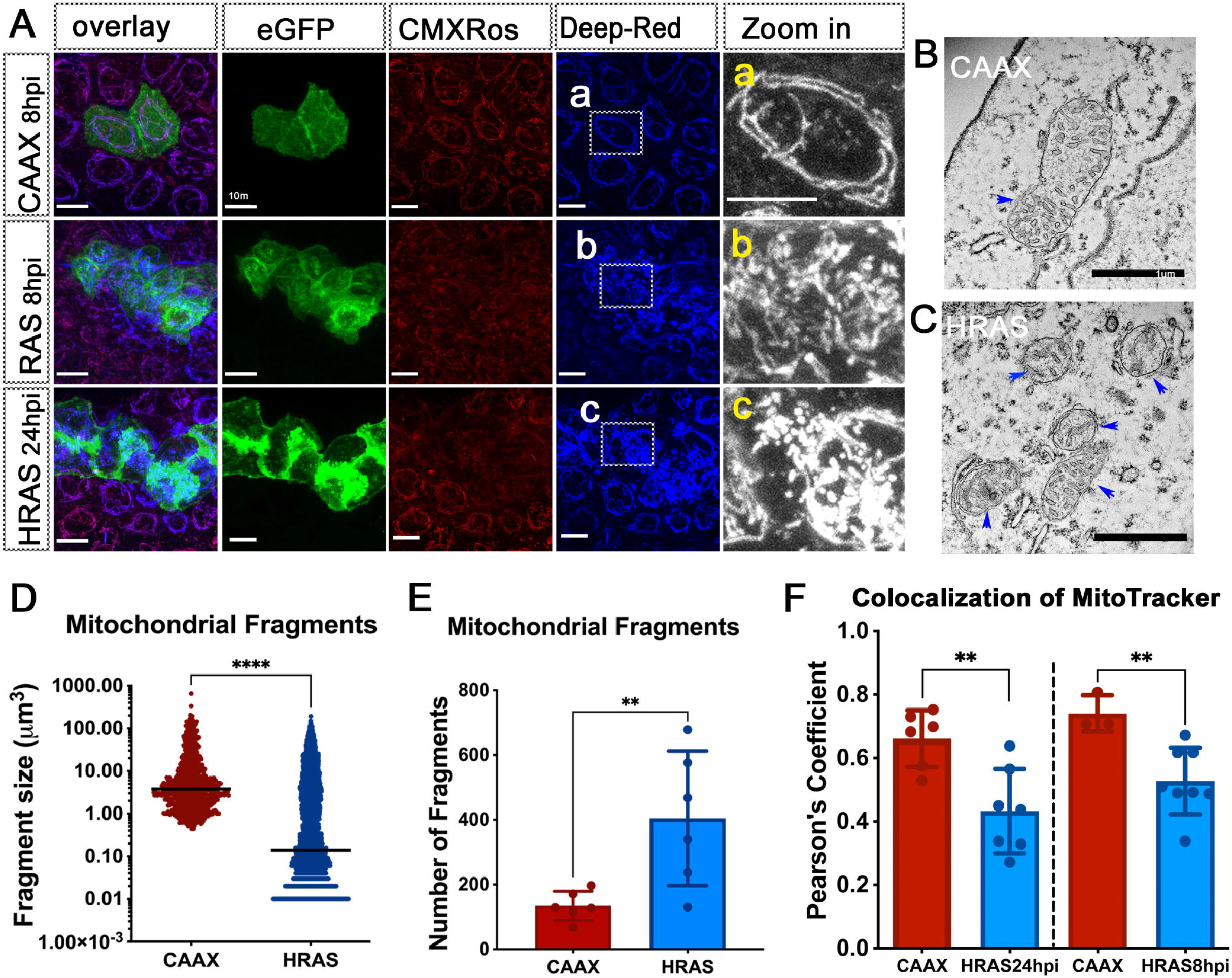
Mitochondria in PNCs are fragmented and have reduced membrane potential. A. Confocal images of mito-trackers CMXRos and Deep-Red stained zebrafish larval skin cells. a, b, c, indicate “Zoom in” area of mito-tracker deep-red image to show details of mitochondrial fragmentation phenotype in HRAS expressing skin PNCs. Scale bar=10μm. B. Electron microscope image of a mitochondrion in normal skin cells of zebrafish larva. C. Electron microscope image of mitochondria in HRAS expressing skin PNC of zebrafish larva. D. Quantification showing increased number of mitochondrial fragments in PNCs (Mann-Whitney test, p=0.0087, n≥8) E. Quantification showing decreased mitochondrial fragment size in PNCs (Mann Whitney test, median, p<0.0001) F. Quantification showing decreased mitochondrial membrane potential in PNCs (unpaired t tests, 24hpi n≥6, p=0.004; 8hpi n≥3, p=0.0099)

**Figure 4.**
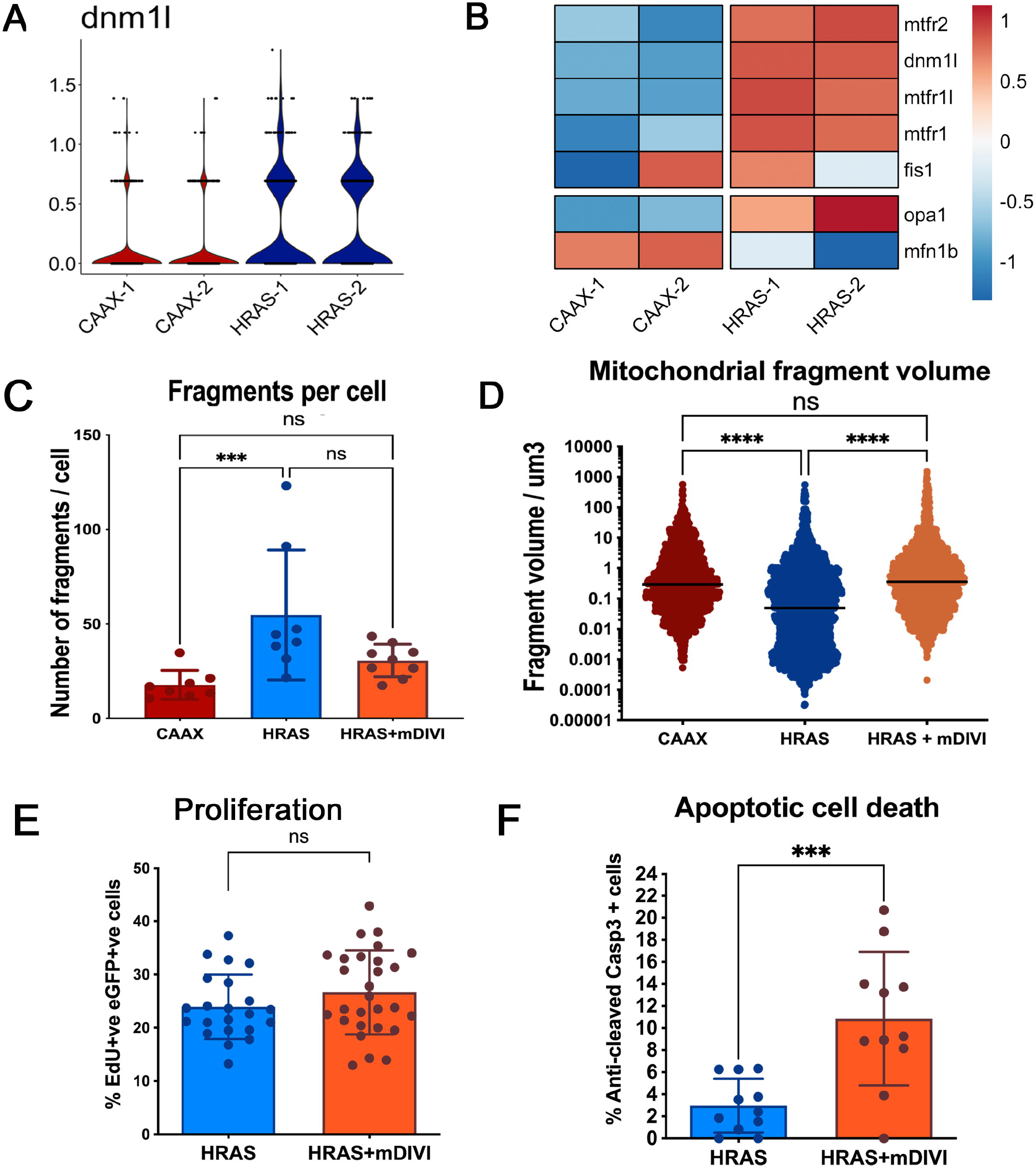
mdivi suppression of Drp1/Dnml1 blocks mitochondrial fission and induces PNC apoptosis. A. Violin plot, depicting normalized expression of *dnml1* in single-cell RNA sequencing dataset, shows that *dnml1* is up-regulated in PNCs vs. CAAX control keratinocytes. B. Pseudobulk differential expression analysis of single-cell RNA sequencing data, showing genes related to mitochondrial fission and fusion in HRAS PNCs vs. control keratinocytes at 24 hpi. Heatmap depicts log fold-change (EdgeR, n=2, FDR < 0.05). C. Quantification showing increased mitochondrial fragmentation can be reversed by mdivi treatment (Kruskal-Wallis test with Dunn’s multiple comparisons, p=0.001, n≥8) D. Quantification showing reduced size of mitochondrial fragments can be reversed by mdivi treatment (Kruskal-Wallis test with Dunn’s multiple comparisons, p<0.0001) E. Quantification showing mdivi does not alter PNC proliferation (unpaired t test, p=0.1889, n≥22). F. quantification showing mdivi induced PNC apoptosis (mean +/− SD, Mann-Whitney test, p=0.001, n=11)

### 3. Mitochondrial fission is an early event in oncogenic HRAS driven preneoplastic cell (PNC) expansion in vivo

Alteration in mitochondrial morphology dynamics was linked to changes of mitochondrial metabolism and was required for transformed cells to drive tumourigenesis in xenograft models^18,27^. To establish whether mitochondrial morphology is changed in PNCs, which might explain altered OXPHOS, we performed *in vivo* live imaging to assess mitochondrial morphology. The vital Mito-Tracker dye revealed that mitochondria became more fragmented in PNCs by 8hpi, the earliest time at which PNCs can be detected by fluorescence and mitochondria remain to be more fragmented at 24hpi (figure 3 A). Compared with stage matched CAAX-EGFP expressing keratinocytes there was a greatly increased number of mitochondrion and the average fragment size was smaller in the PNCs (figure 3 A, D, E). The increased fragmentation of mitochondria was also confirmed by electron-microscopy, in which smaller mitochondria could be seen in the PNCs although we saw no obvious changes to mitochondrial membranes or cristae structure (figure 3 B, C arrowheads).

Increased mitochondrial fission has been linked to mitochondrial damage repair and reduced membrane potential and this could explain the reduced mitochondrial respiratory reserve that we detected ^28,29^. We took advantage of two MitoTracker dyes with different sensitivities to mitochondrial membrane potential^30^. While the MitoTracker-DeepRedFM is insensitive to mitochondrial membrane potential and labelled each mitochondrion in its entirety (figure 3 A, a, b, c), the mitochondrial membrane potential dependent dye, MitoTracker Red CMXRos, was much fainter and occasionally missing from mitochondria in PNCs compared to MitoTracker-DeepRedFM (figure 3 A). Co-localization analysis of the two MitoTrackers showed a reduced membrane potential in PNCs as early as 8hpi (figure 3 A, F) and this reduction was maintained at 24hpi (figure 3 A, F) corroborating reduced mitochondrial maximum respiration that we observed.

### 4. Drp1/Dnml1 inhibitor mdivi blocks mitochondrial fragmentation and induces PNC apoptosis

It has previously been reported previously that the small GTPase Drp1 (called dnml1 in zebrafish) drives mitochondria fragmentation in cancer cells carrying oncogenic RAS mutations in vitro ^18,29^. Using previously generated single-cell RNA sequencing data (Elliot et al. unpublished) we found that *dnml1* was indeed up-regulated in PNCs at 24hpi (figure 4 A). Alongside *dnml1* we also saw upregulation of other mitochondrial fission proteins such as *fis1, mtfr1, mtfr1l, mtfr2*^31 32^(figure 4B). Conversely, one of the key mitochondrial outer membrane fusion protein *mfn1b* is down regulated ^17^(figure 4 B). Thus, gene expression changes amongst mitochondrial dynamic regulators support enhanced mitochondrial fission that we saw. The increased mitochondrial fission has been shown to be required for RAS mutant tumour formation in a xenograft model^18,29^. Therefore, we speculated that mitochondrial fission might be necessary for PNC initial development. To test whether reversing the mitochondrial fission phenotype might negatively impact on PNC development, we utilized a Drp1/Dnml1 inhibitor mdivi, which has been used in multiple models to reverse Drp1/Dnml1 mediated mitochondrial fission phenotype ^33,34^. Due to its ability of reduce pathological mitochondrial fission and protection from cell death in degenerative diseases, mdivi holds promise in the treatment of neuro-degenerative diseases and is safe for human use^34^. Mdivi has also been shown to enhance cancer cell death through inhibiting Drp1^35^. In zebrafish, mdivi has previously been show to protect hair cells from cisplatin induced death^36^. When we treated PNC bearing larvae with mdivi, we could reduce the mitochondrial fragmentation of PNC in vivo within 2 hours of treatment (Figure 4 B, C, D).

Using mdivi treatment to reverse mitochondrial fragmentation, we found that PNC proliferation remained unaltered (figure 4 E), but we did detect significantly increased apoptotic cell death of PNCs (figure 4 F). These data suggest that oncogenic RAS expressing PNCs undergo metabolic adaption through Dnml1 mediated mitochondrial fission and that this is required for PNC survival. Moreover, we show how this could be efficiently targeted by small molecule mdivi for therapeutic PNC elimination.

In summary, our studies have established an enabling role for rapid metabolic adaption in cells that switch on RAS mediated oncogenic pathways, by promoting their survival and expansion in vivo at the preneoplastic stage. We show that “Warburg metabolism” is required for PNC hyper-proliferation and increased availability of glucose in vivo further promotes PNC proliferation, and which in turn maximises their expansion. We also find that a rapid change in mitochondrial dynamics and function, is required for PNC survival during their rapid proliferative expansion. The excess glucose resource mediated PNC hyper-proliferation might provide a mechanism linking dietary high sugar intake to cancer incidence that has been observed in humans ^37,38^. Importantly, we provide evidence that the metabolic adaptation pathways might be targeted to selectively eliminate PNC through re-purposing currently available drugs, thus might provide effective tumour prevention strategies for human with RAS driven cancer predisposition syndromes^39^.

## Materials and Methods

### Zebrafish maintenance and breeding

Adult zebrafish were maintained in the Bioresearch & Veterinary Services (BVS) Aquatic facility in the Queen’s Medical Research Institute, The University of Edinburgh. Housing conditions were described in the Zebrafish Book (Westerfield, 2000) with 14/10 hours light/dark cycle and water temperature of 28.5 °C. Adult K19 Gal4 or K4 Gal4 driver fish were set for pair mating with either UAS:RAS or UAS:CAAX fish in pair mating tanks. Next morning, the fertilized embryos were collected within 2 hours of removing dividers to ensure synchronous development. The embryos were maintained in 90 mm petri dishes (maximum of 50 embryos/dish) containing 0.3× Danieau’s solution (17.40 mM NaCl, 0.21 mM KCl, 0.12 mM MgSO_4_•7H_2_O, 0.18 mM Ca(NO_3_)_2_, 1.5 mM HEPES), in a 28.5 °C incubator. All experiments were conducted with local ethical approval from The University of Edinburgh and in accordance with UK Home Office regulations (Guidance on the Operations of Animals, Scientific Procedures Act, 1986).

### Transgenic Zebrafish strains

The zebrafish lines used in this study include Tg(krtt1c19e::KalTA4-ER^T2^; cmlc2::eGFP)^ed201^ (hereafter K19 Gal4 driver) primarily drives basal keratinocytes. Tg(krt4::KalTA4-ER^T2^; cmlc2::eGFP)^ed202^ (hereafter K4 Gal4 driver) primarily expressed in superficial keratinocytes^40^ Tg(UAS::eGFP-HRAS^G12V^; cmlc2::eGFP)^ed203^ (hereafter UAS:RAS)^11^. Tg(UAS::eGFP-CAAX; cmlc2::eGFP)^ed204^ (hereafter UAS:CAAX)^11^

### Zebrafish skin Preneoplastic cell induction and drug treatment

The detailed induction protocol has been described previously^10^. In brief, the induction solution consisted of 0.3× Danieau’s solution with 5 μM 4-OHT and 0.5% DMSO to enhance penetration of 4-OHT. A 10 mM stock of 4-OHT dissolved in 96% ethanol was stored at −20 °C protected from light. At 48 hours post-fertilisation (hpf), selection marker positive (green heart) larvae were transferred to petri dishes containing 20 ml 4-OHT induction solution, at 50 larvae per dish. The larvae were then maintained at 28.5 °C in the dark. For Oxygen consumption rate (OCR) and glycolysis analysis using Seahorse ^®^ analyser, the larvae were anaesthetised with 55 mg/L eugenol (Sigma) at 22 hours post-induction (hpi), and screened. From the same clutch, larvae with GFP positive skin cells were collected as pre-neoplastic cell (PNC) group, and larvae with GFP negative skin were collected as Wild Type (WT) siblings group. For drug treatment, larvae were screened at 8hpi, and positive larvae from CAAX group were used as controls. The sorted embryos were placed in fresh 4-OHT induction solution with appropriate drugs.

### Zebrafish larvae wholemount Seahorse ^®^ Flux analysis for oxygen consumption rate (OCR) and extra cellular acidification rate (ECAR)

Oxygen consumption rates (OCR) and extracellular acidification rates (ECAR) were measured using the Seahorse XFe24 Extracellular Flux Analyser (Agilent Technologies). 4-OHT treated embryos at 24 hpi were anaesthetised by treating with 55 mg/L eugenol (Sigma), screened, and loaded into an islet capture plate (Agilent Technologies) (1 embryo/well) in a randomised order. Islet capture screens were added to keep embryos in position. A final volume of 525 μl 0.3× Danieau’s Solution was added to each well. The plate was then loaded into the XFe24 analyser. Experiments were carried out at approximately 27 °C (+/−1 °C). Reagents used for manipulating glycolysis and respiration were obtained and used at final concentrations as follows; 2-Deoxyglucose (100mM) was used for inhibiting glycolysis (and associated ECAR) and was obtained from the Seahorse XF Glycolytic Rate Assay Kit (Agilent Technologies 103344-100. FCCP (5μM) was used to reveal maximal respiratory capacity (and associated OCR) by uncoupling mitochondrial respiration. Antimycin/Rotenone (2μM/2μM) was used to abolish respiration (for subtraction of non-respiratory OCR). FCCP, Antimycin and Rotenone were obtained from the Seahorse XF Cell Mito Stress Test Kit (Agilent technologies 103015-100). Note: Oligomycin were tested but fail to penetrate zebrafish larvae and did not provide any meaningful reading. For calculating glycolytic flux, baseline ECAR was measured six times prior to addition of 2-deoxyglucose. Glycolytic ECAR was revealed by the drop in ECAR subsequent to the addition of 2-deoxyglucose. In the glycolysis test, FCCP was added two measurements after, to estimate maximal respiration using the parallel OCR measurements. ECAR rise post 2-deoxyglucose and FCCP treatments are not considered glycolytic flux, and reflect other acidification phenomena including respiratory CO_2_ production.

For focused respiration tests, four baseline OCR measurements were taken, prior to addition of FCCP. Four measurements were taken after FCCP, prior to addition of Antimycin and Rotenone. A further eight measurements were taken after respiratory inhibition to allow time for OCR to drop. Maximal respiration was calculated by subtracting the Antimycin/Rotenone inhibited OCR from the peak FCCP stimulated OCR (after FCCP). Post run analysis was completed using Agilent Seahorse Wave software (Version2.6) and Prism.

### In vivo MitoTracker staining and imaging

4-OHT treated larvae were selected for positive signal in skin cells at either 8hpi or 24hpi and then incubated with 500 nM MitoTracker Red CMXRos (Thermo Fisher Scientific M7512) and 500 nM MitoTracker Deep Red FM (Thermos Fisher Scientific M22426) in 1.5ml Eppendorf tubes for 30 minutes at 28.5 °C in the dark followed by washing 3 times with 0.3× Danieau’s solution for 30 seconds. For mDIVI treatment experiments, embryos were treated two hours prior to staining. After washing, larvae were immediately embedded in 0.8% low melting temperature agarose in glass-bottom dishes for confocal imaging.

### Zebrafish larval EdU (5-ethynyl-2⍰-deoxyuridine, a nucleoside analog of thymidine) incorporation and staining

4-OHT induced larvae were selected for positive signal at 8hpi, and if required appropriate drugs were added to fresh 4-OHT induction solution for further treatment. 2nl EdU solution (10mM) was injected into the yolk of individual larva and after 2.5 hours incubation, if required 24hpi, larvae were fixed with 4% PFA for 30 minutes at room temperature. EdU staining was carried out using the Click-iTTM Plus Edu Alexa FluorTM 647 Imaging Kit (Thermo Fisher Scientific, C10640) following manufacturers instructions. In brief, larvae were washed thrice in PBS containing 0.5% Triton X-100 (PBST) for 5 minutes, and blocked with PBST containing 3% (w/v) Bovine Serum Albumin (Sigma-Aldrich, Gillingham, UK) for 1 h. This was followed by 30 min incubation with the Click-it Plus reaction cocktail using 250 μl of reaction cocktail per 10 larvae. Following EdU labelling, larvae were washed thrice with PBST for 5 min and re-blocked with PBST containing 5% (v/v) Goat Serum (Sigma-Aldrich, Gillingham, UK) for 2 h. eGFP immunostaining was performed with rabbit monoclonal anti-GFP antibody (1:200; cat. 2956, Cell Signaling Technology, London, UK) and Alexa Fluor 488 Goat anti-Rabbit secondary antibody (1:250; A-11008, Invitrogen, Fisher Scientific, Loughborough, UK) as described (van den Berg et al., 2019). Stained larvae were stored at 4 °C in a glycerol based antifadent mountant (AF1, CitiFluor, Hatfield, PA, USA) until mounted for imaging.

### Wholemount immunostaining for activated-caspase-3

Larvae were fixed in 4% PFA for 2 hours at room temperature followed by incubation in MeOH overnight at −20 °C. Larvae were gradually re-hydrated at room temperature before washing thrice with 0.1% PBST for 5 minutes and then incubated in blocking solution for 2 hours with gentle shaking. This was followed by overnight incubation with the primary antibody (Purified Rabbit Anti-active caspase 3 (BD 559565) 1:200 in Block solution) at +4°C with shaking. The following day, larvae were washed 6 x 20 min with 0.1%PBT at room temp with shaking, followed by incubation for 2 hours in secondary antibody (Alexa Fluor 633 goat anti-rabbit IgG (H+L) (Invitrogen A21071) 1:250 in block solution). The larvae were washed 3 × 15 min in 0.1% PBT at room temperature. Following staining, larvae were stored in Citi Fluor at +4°C before mounting for confocal imaging.

### Confocal imaging

All images for fixed samples were acquired with Leica confocal SP8, using HCX PL APO 40× water immersion objective lens. Live imaging for MitoTracker were carried out using HCX PL APO 63× glycerol immersion objective lens. Appropriate lasers and collection gates were selected according to excitation and emission wavelength for each fluorophore.

### Image data analysis

All image data analyses were carried out using IMARIS 8.2 or IMARIS 9.0. Percentage of EdU or active caspase-3 positive cells within eGFP positive cells were manually counted. Images from MitoTracker Deep Red FM staining channel were used for mitochondrial morphology assessments. First, isolation of PNCs or CAAX clones in GFP channel using the “Surfaces” feature. Mask MitoTracker channel outside “Surface” to generate a MitoTracker channel specific within GFP^+^ cells. Isolation of mitochondrial fragments using the “Surfaces” feature based on absolute intensity of the masked MitoTracker channel. Use of resulting Surface statistics to report mitochondrial fragment number and size.

### Statistical analysis

Statistical analysis was done using Prism6 (GraphPad). ECAR and OCR trace data were analysed using Two-way ANOVA with Sidak multiple comparison tests at each measuring cycle. ECAR measurements after 2-DG addition and OCR measurements after FCCP addition were also analysed using unpaired t test. For all other graphs, when comparing two groups unpaired *t* tests were used for samples with equal SD or Mann Whitney test was used for samples with different SD. For mitochondrial fragments and volume comparison this included the CAAX, HRAS and HRAS+mdivi groups, Kruskal-Wallis test with Dunn’s multiple comparisons were performed.

### Gene expression analysis

Single-cell RNA sequencing data from Elliot et al. was re-analysed to obtain a comparison of HRAS expressing PNCs and CAAX expressing control basal keratinocytes. Cells from 24hpi samples labelled “Basal Keratinocyte” or “PNC” were grouped by sample and their raw counts were summed across cells to represent “pseudobulk” RNA sequencing samples. Normalisation and differential expression analysis (glmQLFitTest) were carried out using edgeR ^41,42,43^. Gene-set enrichment analysis ^44,45^ was carried out upon the differential expression results using gene-sets obtained from the *Danio rerio* (26246) Gene Ontology database^45,46^.

## Supplemental Material

Supplemental figure 1

A. Trace annotation for Oxygen Consumption Rate measurement from Seahorse XF^®^ Analyser, for data showed in main figure 2 A, B.

Supplemental figure 2

A. Graph shows quantification for anti-active-caspase 3 staining in CAAX cells from larvae treated with mDIVI, metformin, Glucose, showing CAAX cells are not affected by the treatments. (One-way ANOVA analysis with Dunnet’s multiple comparison, n≥9) B. Graph shows quantification for EdU positive CAAX cells in glucose treated larvae compared with untreated control and CAAX cells are not affected. Mann-Whitney test, n≥14, p>0.99. C Graph shows quantification for EdU positive CAAX cells in mdivi treated larvae compared with untreated control and CAAX cells are not affected. Mann-Whitney test, n≥14, p=0.1933. D. Graph shows quantification for EdU positive CAAX cells in metformin treated larvae compared with untreated control and CAAX cells are not affected. Unpaired *t* test, n≥11, p=0.3957. E. Graph shows quantification for EdU positive CAAX cells in lonidamine treated larvae compared with untreated control and CAAX cells are not affected. Unpaired *t* test, n≥12, p=0.1169.

## Acknowledgement

The authors thank the technical support from the Confocal Advanced Light Microscopy, and the BVS aquatic facility units at the University of Edinburgh. The authors acknowledge fundings from Wellcome Trust Sir Henry Dale Fellowship to Y.F. (100104/Z/12/Z); Cancer Research UK Early Detection Award to Y.F. (C38363/A26931); A.M.E. was supported by a Medical Research Council PhD Studentship (MR/N013166/1); L.K was funded by university of Edinburgh principle fellowship and Wellcome Trust Beit Prize to Y. F.; H. M. was supported by a postdoctoral fellowship from the Sigrid Jusélius Foundation. R.N.C was funded by a Wellcome Trust New Investigator award 100981/Z/13/Z to N.M.M.

## Notes

### Competing Interest Statement

The authors have declared no competing interest.

