## Supplementary figures and images for "Preneoplastic cells switch to Warburg metabolism from their inception exposing multiple vulnerabilities for targeted elimination"

### supplemental figure 1

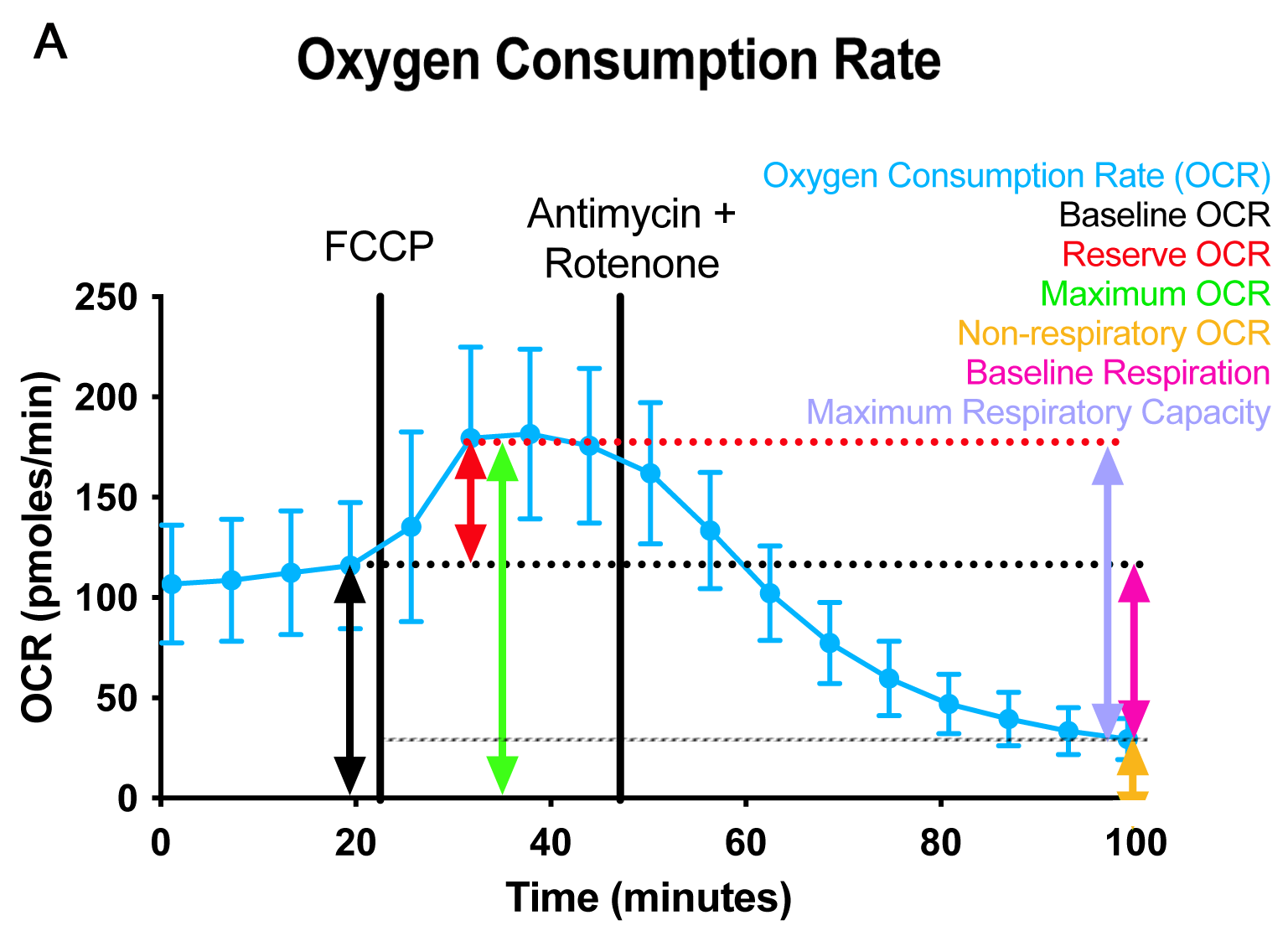

### supplemental figure 2

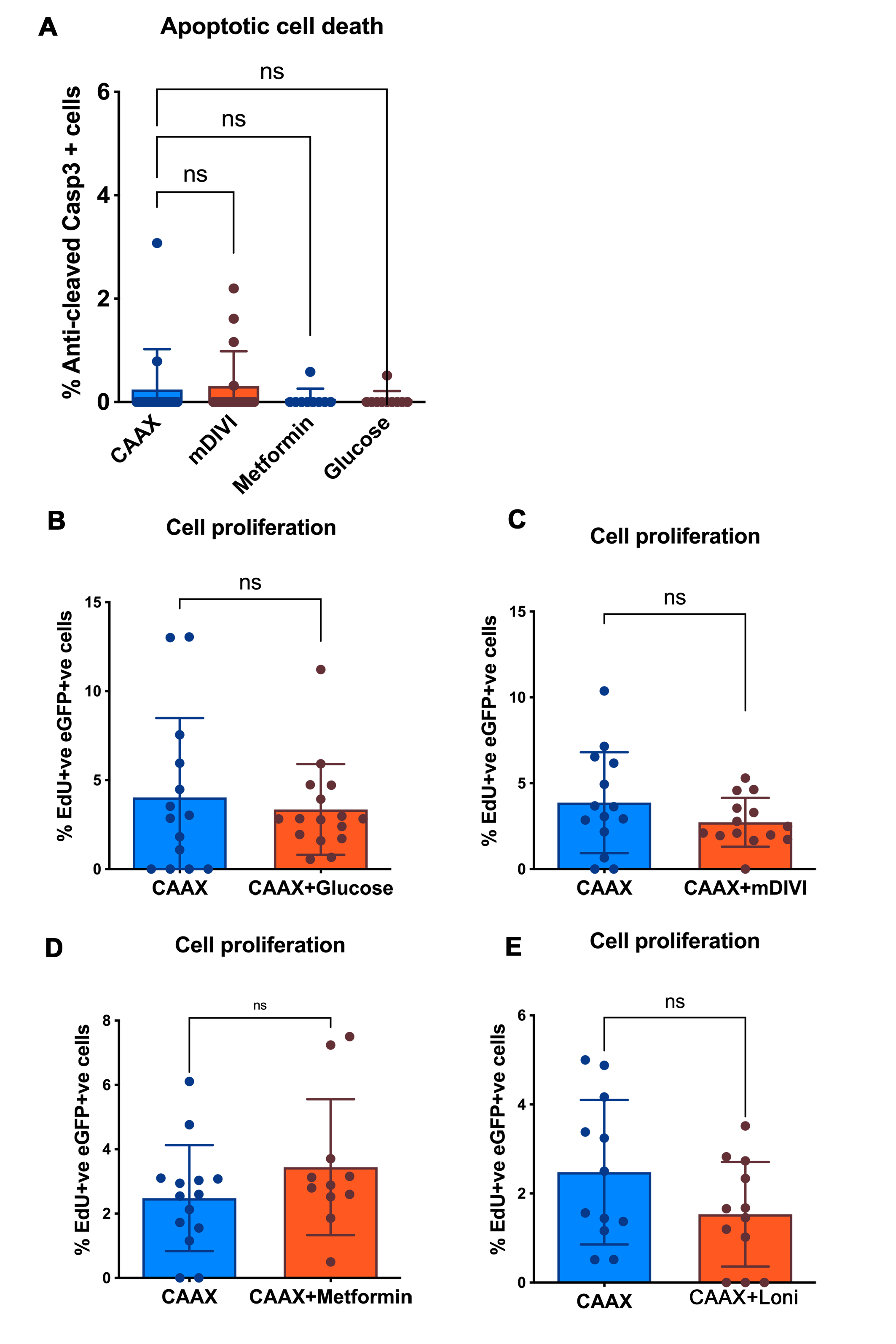
